# QuantGUV: Quantifying Encapsulation Efficiency of Small Molecules in GUVs

**DOI:** 10.1101/2025.10.23.684161

**Authors:** Zak Marshall, Reshma Bano, Pasha Dylan, Luisa Trifan, Callum Mckeaveney, AndréP. Gerber, Wooli Bae

## Abstract

Synthetic cells, constructed through the self-assembly of small molecules, are designed to mimic life-like behaviours by encapsulating functional molecules. For such synthetic cells to accurately replicate cellular reactions, it is critical that the concentrations of encapsulated molecules mirror those in living systems, as reaction kinetics and cellular network states are highly sensitive to these concentrations. However, methods for precisely determining encapsulation efficiency in synthetic cells at single cell resolution have been limited. To address this challenge, we developed QuantGUV, a software-driven, image-based analysis method that determines the concentrations of fluorescent molecules encapsulated within giant unilamellar vesicles (GUVs). We used QuantGUV to measure the encapsulation efficiencies of fluorescent molecules, ranging in size from 0.5 nm to 20 nm. These measurements were conducted on GUVs formed via the water-in-oil emulsion transfer method under various experimental conditions. Using QuantGUV, we have measured the encapsulation efficiencies of three fluorescence molecules, sulforhodamine B, mEGFP and polystyrene bead, in GUVs formed via the water-in-oil emulsion transfer method. The encapsulation efficiencies for polystyrene bead was close to 100% in most of the conditions while sulforhodamine B and mEGFP’s encapsulation efficiencies depended on the parameters during the GUV formation such as concentrations of lipids and oil-water ratio during the GUV formation. By providing crucial insights into encapsulation efficiencies, QuantGUV offers a valuable tool to monitor the building quantitative synthetic cell systems with accurately controlled internal environments which is a critical step towards the creation of synthetic cells.

## 2 Introduction

A hallmark of living organisms is compartmentalisation, a fundamental principle in which a membrane physically and functionally separates a living system from its surroundings [1, 2]. Fuelled by recent advances in membrane biophysics, polymer physics, and bottom-up synthetic biology, it is now possible to construct artificial compartments with various physical properties such as size, curvature and fluidity to mimic different properties of biological systems [3–7]. Giant unilamellar vesicles (GUVs) are one of the key systems for studying compartmentalisation, as they resemble the size of living cells and can be produced at high amounts [8–16]. The GUVs can encapsulate functional molecules, including fluorescence compounds or a cell-free transcription-translation (TXTL) system to form synthetic cells that replicate core biological processes like information processing, membrane dynamics [17–24], molecular transport [25–28], and also developing possible drug delivery systems [29–31].

To correctly mimic biochemical reactions, it is essential to encapsulate functional molecules at desired concentrations as the rates and equilibrium dynamics depends on them. However, a lot of currently available methods to form GUVs are subjective to potential leakage and encapsulation concentration heterogeneities [32–38](Figure 1a). A limitation of these methods is their inability to guarantee the retention of initial molecular concentrations following GUV formation. Although it is straightforward to compare the relative encapsulation efficiency of different methods or compartments, this information is not sufficient to build predictable synthetic cells. Additionally, the overall yield of multi-step enzymatic reactions, such as the in vitro synthesis of proteins, decreases exponentially with the number of steps involved [39, 40].

**Figure 1:**
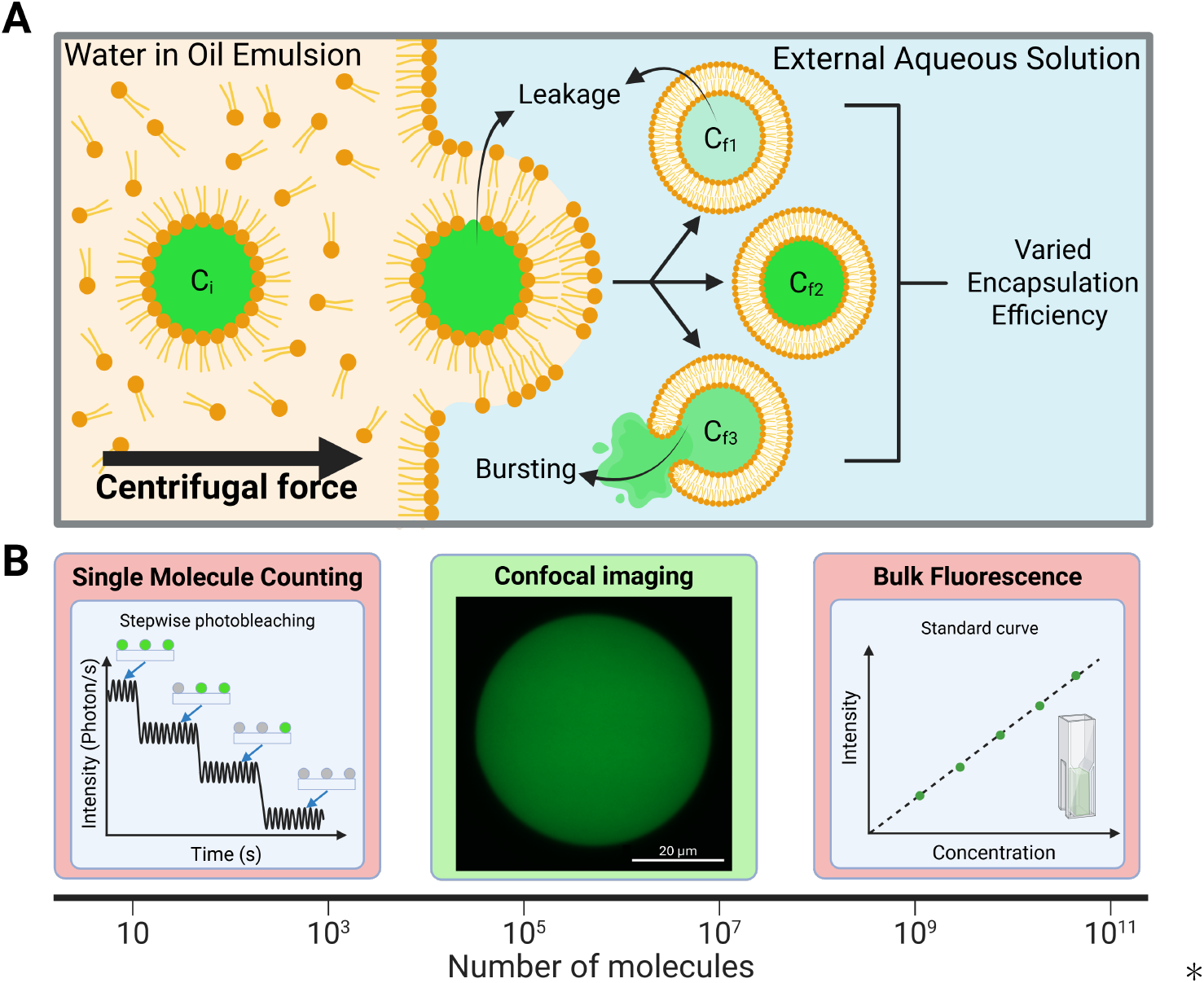
Challenges in encapsulation of molecules and determining the encapsulation efficiency A. Schematic of GUV formation using inverted emulsion transfer method displaying that the initial concentration, C_i_, can change due to Leakage, C_f1_, random encapsulation, C_f2_ and bursting, C_f3_. B. Methods for quantifying the concentration of fluorescence molecules, Left red, single molecule counting, Right red, Bulk fluorescence using standard curves. Central green, confocal microscopy with reduced background. Created in BioRender. Marshall, Z. (2025) https://BioRender.com/qa7p0vv

Therefore, there have been several attempts to quantify or enhance encapsulation of functional molecules in GUVs. Sun et.al described a method of photolysis and single-molecule detection using confocal imaging to quantify the dyes encapsulated. Encapsulated fluorescent dyes were released by laser lysis of GUVs and the measured fluorescence was fitted to a diffusion model to estimate encapsulation efficiency [41]. They reported an encapsulation efficiency of 36.3 ± 18.9 % for unilamellar / oligo lamellar vesicles. Matosevic et al. used a microfluidic device to produce GUVs, and by comparing the measured fluorescence intensity of the droplets pre and post transfer, they were able to report an average encapsulation efficiency of 83 % [42]. Göpfrich et al. reported a method for producing GUVs using charged lipids in a “one-pot” method. This method was reported to retain high encapsulation efficiency, however was not quantified [43]. Liu et al. used colourimetry with the Bradford method for quantifying protein encapsulation efficiency in the supernatant of GUVs formed from the emulsion transfer method [44] and reported an average of 97.75 ± 1.73 % for enzymes such as RNA Polymerase. Recently, Supramaniam et al. developed a microfluidic analysis chamber through which vesicles formed through inverted emulsion phase transfer were captured in an analysis chamber and lysed by both chemical and optical methods [45]. By using single molecule microscopy across a microarray they counted spots and extrapolated to a standard curve to give absolute quantification with a mean encapsulation efficiency of 11.4 %.

However, the examples above only give average values or involve specialised equipment and procedures with relatively high barrier to entry, such as microfluidic techniques or single-molecule fluorescence microscopy. Current techniques are unsuitable for widespread adoption because they rely on specialised, high-barrier-to-entry equipment or bulk measurement methods which fail to address the intrinsic heterogeneity of GUVs size and content. While quantification at a single vesicle level with high throughput is desired, the physical scale of the GUVs prevent the use of established methods such as single molecule counting and obtaining a calibration curve from bulk solutions (Figure 1b). Developing a standardised high-throughput approach to quantify encapsulation efficiency with a low barrier for entry of different sorts of molecules would enable wider adoption of GUVs for engineering and synthetic biological purposes and enable more precise modelling of experimental results with mathematical modelling.

Hereom, we present QuantGUV, a simple standardised, software-driven pipeline for a high-throughput quantification of concentrations of encapsulated fluorescent molecules within GUVs (figure 2). In brief, we generated standard curves of the intensities of fluorescence molecules at different concentrates inside of a GUV. Specifically, we took images of bulk dye solution and subtracted background intensities generated from images of empty GUVs in bulk dye solutions. After taking images the encapsulated molecules of interest in GUVs, an automated process determines the concentrations of the molecules by interpolating the estimated concentration from the standard curve. We produced GUVs using the water-in-oil emulsion transfer method, one of the most-widely used method with applications for encapsulation of molecules of interest [8, 46]. Encapsulation efficiency was calculated for each individual GUV as the ratio of the encapsulate concentration within the GUV to the total encapsulate concentration initially added to the emulsion phase, expressed as a percentage. We used QuantGUV to characterise encapsulation efficiencies of three molecules with different sizes - Sulforhodamine B (559 Da), monomeric enhanced green fluorescent protein (mEGFP) (27 kDa) and fluorescent spheres (2.6 MDa) at different conditions during the GUV formation.

**Figure 2:**
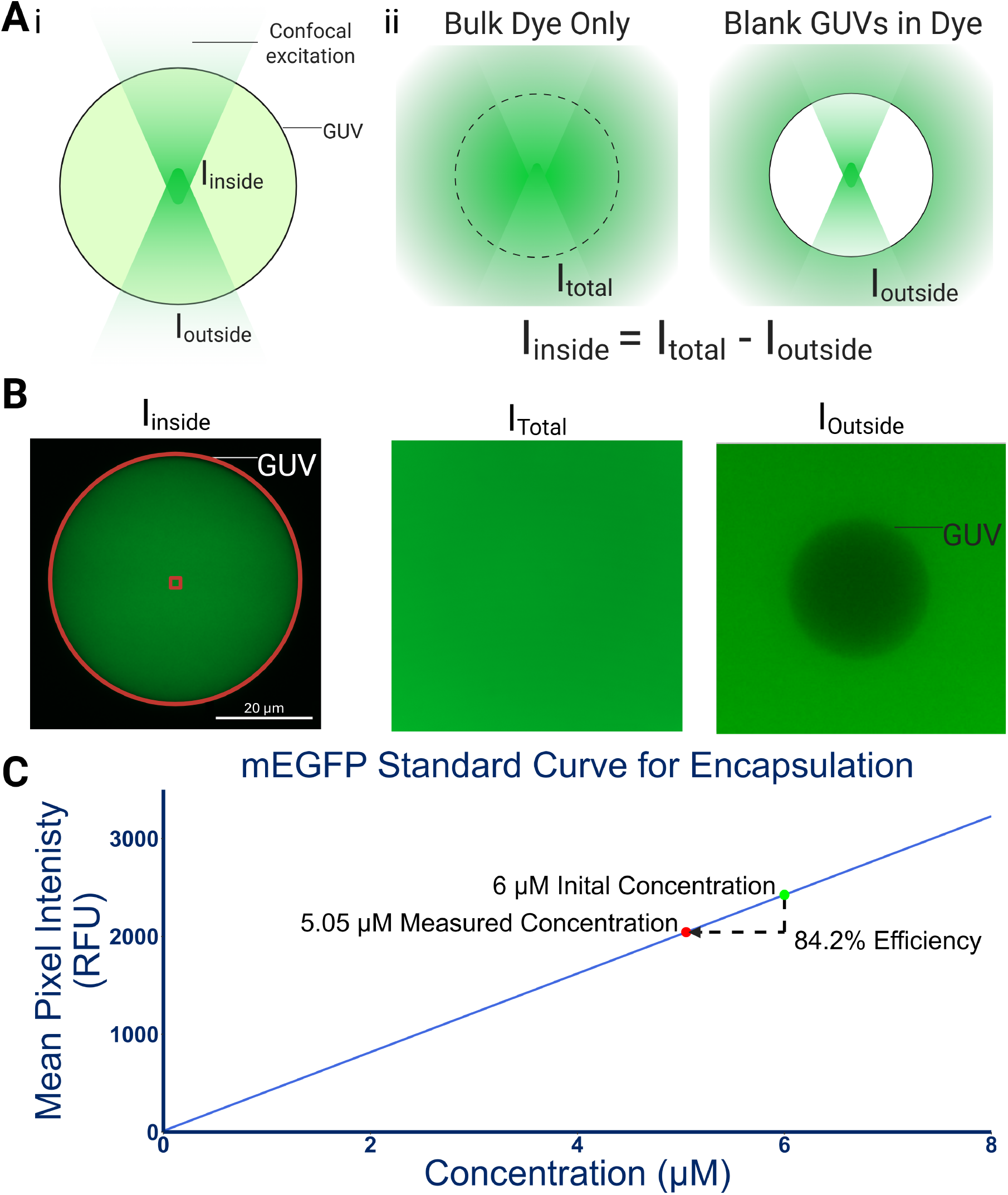
QuantGUV calibration curves A) A schematic illustrating how the bright background correction works. To correctly estimate I_Inside_, I_Outside_ and I_Total_ are measured separately using empty GUVs in bulk solutions and bulk solutions only B) Confocal images demonstrating the working principle. From left to right, a GUV encapsulating mEGFP (I_Inside_), a bulk mEGFP solution (I_Total_) and a blank GUV within a solution containing mEGFP (I_Outside_), The image contrast and highlights have been enhanced for ease of viewing. C) mEGFP standard curve correlating known protein concentration with mean pixel intensity (RFU) calculated from I_Total_ - I_Outside_. The equation of the line is used to set the RFU for initial concentration of protein in the solution (green dot) the same equation is used to interpolate the I_Inside_ of the GUV above and determine internal concentration (red dot). Encapsulation efficiency is determined by 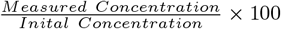× 100. Created in BioRender. Marshall, Z. (2025) https://BioRender.com/953ins9

## 3 Results

### 3.1 QuantGUV Standard Curve Generation

As the number of encapsulated molecules inside of a GUV with a diameter of 10 µm at 1 µM is over 300,000, it is not possible to count individual molecules at this range. Therefore, we can only obtain fluorescence signal from a point inside of a GUV using confocal microscopy which comes from encapsulated molecules. While this signal should be proportional to the concentration of encapsulated molecules, it is not possible to directly infer the concentration of molecules directly from the signal as fluorescence signal depends on various factors including the intensity of laser, numerical aperture of objective lens and quantum yield of detectors. Environmental dependency of fluorescence quantum yield makes this task even more challenging.

Therefore, we need a standard curve which can relate the internal fluorescence signal of GUVs to a concentration of the fluorescence molecule (Figure 2, left, *I*_*Inside*_). However, taking the fluorescence signal of bulk solution (Figure 2, middle,*I*_*total*_), and relate it with the fluorescence signal measured inside of the GUVs would not work. Even when using confocal microscopy, out-of-focus fluorescence signals will not be zero. This is represented by *I*_*total*_ = *I*_*Inside*_ + *I*_*Outside*_ with *I*_*outside*_ denoting fluorescence signals coming from outside of GUV volume. While it would be ideal to use fluorescence signals from GUVs bearing defined concentrations of fluorescent molecules as standards, it is not straightforward to make such GUVs due to the non-ideal encapsulation efficiencies and heterogeneity of the GUVs, which was the original problem that we are dealing with. Although it is not possible to measure.*I*_*Inside*_ directly, it is possible to measure *I*_*outside*_ and *I*_*total*_ directly and work out the *I*_*Inside*_ as *I*_*Inside*_ = *I*_*Total*_*− I*_*Outside*_. We hypothesised that the *I*_*Outside*_, can be measured by placing blank GUVs in solution with different concentrations of fluorescence molecules outside of the GUVs (Figure 2a, right).

We have experimentally measured *I*_*Total*_ and *I*_*Outside*_ for each fluorescent molecule investigated at different concentrations. To measure the *I*_*Total*_, a dilution series of the molecule of interest was made in internal aqueous solution (IAS) and placed into a 96 well microplate well and imaged on a confocal plater reader. The mean pixel average across 10 images is assigned to *I*_*Total*_. Next, we prepared empty GUVs and likewise placed them into the wells and added fluorescence molecules to the outside of the GUVs. QuantGUV is then used to detect the fluorescence voids from empty GUVs against the bright fluorescent background. The fluorescence voids are due to the lack of fluorescence in the vesicle lumen, however they still display an intensity, which is the *I*_*Outside*_ component. The subtraction of this value from the *I*_*Total*_ concentration was used to calculate intensity values and generation of a standard curve as shown in figure 2b (for mEGFP).

### 3.2 QuantGUV Quantification of GUVs

As the accurate measurement of the *I*_*Total*_, *I*_*Inside*_ and *I*_*Outside*_ is important in obtaining a good calibration curve, we have developed the QuantGUV pipeline to correct for the errors coming during the imaging process. Firstly, to minimise the effect from the lipid membranes, which could affect the quantum yield of fluorescence molecules, QuantGUV determines the geometric centre of the detected GUVs and measure the fluorescence signal from that region. Secondly, GUVs are immobilised on surface for stable signal acquisition. Also, a series of computational algorithms are applied to excludes any out-of-focus vesicles and correct for background fluorescence (See Methods for details). After the quantification of images, the intensity values were normalised by dividing the intensity by the integration time to match with the standard curve. This allows for wider dynamic range and higher signal to noise value for improved data quality. After determining its concentration, it is compared to the initial concentration of molecules in the IAS. The vesicle detected in figure 2b, left, was normalized and plotted on the standard curve in figure 2c where the concentration detected and initial concentration can be compared. In the initial emulsion 6 µM of mEGFP was used and by using the standard curve the estimated concentration within the vesicle was 5.05 µM which is a 84.2% encapsulation efficiency.

### 3.3 Quantifying relevant molecular sized molecules in GUVs

To represent a typical biochemical reaction involving molecules with different sizes, we measured encapsulation efficiencies of Sulforhodamine B (SRB, 1 nm), monomeric enhanced green fluorescent protein (mEGFP, 4 nm) and FluoSpheres (FS, 20 nm). SRB is a small fluorescent dye 580 Da, with a size similar to nucleotides or short peptides. mEGFP is a fluorescent polypeptide 27 kDa, with a size similar to small enzymes or protein subunits. FS are styrene spheres modified with a fluorophore 2.6 M Da, the FS is a similar size to a bacterial ribosome and other large supremolecular complexes.

We have tested 5 % IAS ratio with varying concentrations of lipid within the Lipid in Oil (LiO) mixture to see if a significant difference in encapsulation could be achieved. At all three LiO concentrations, the vesicles were successfully formed and encapsulated the fluorescent molecules (Figure 3A). Thereby, data was screened to remove out of focus GUVs with each condition.

**Figure 3:**
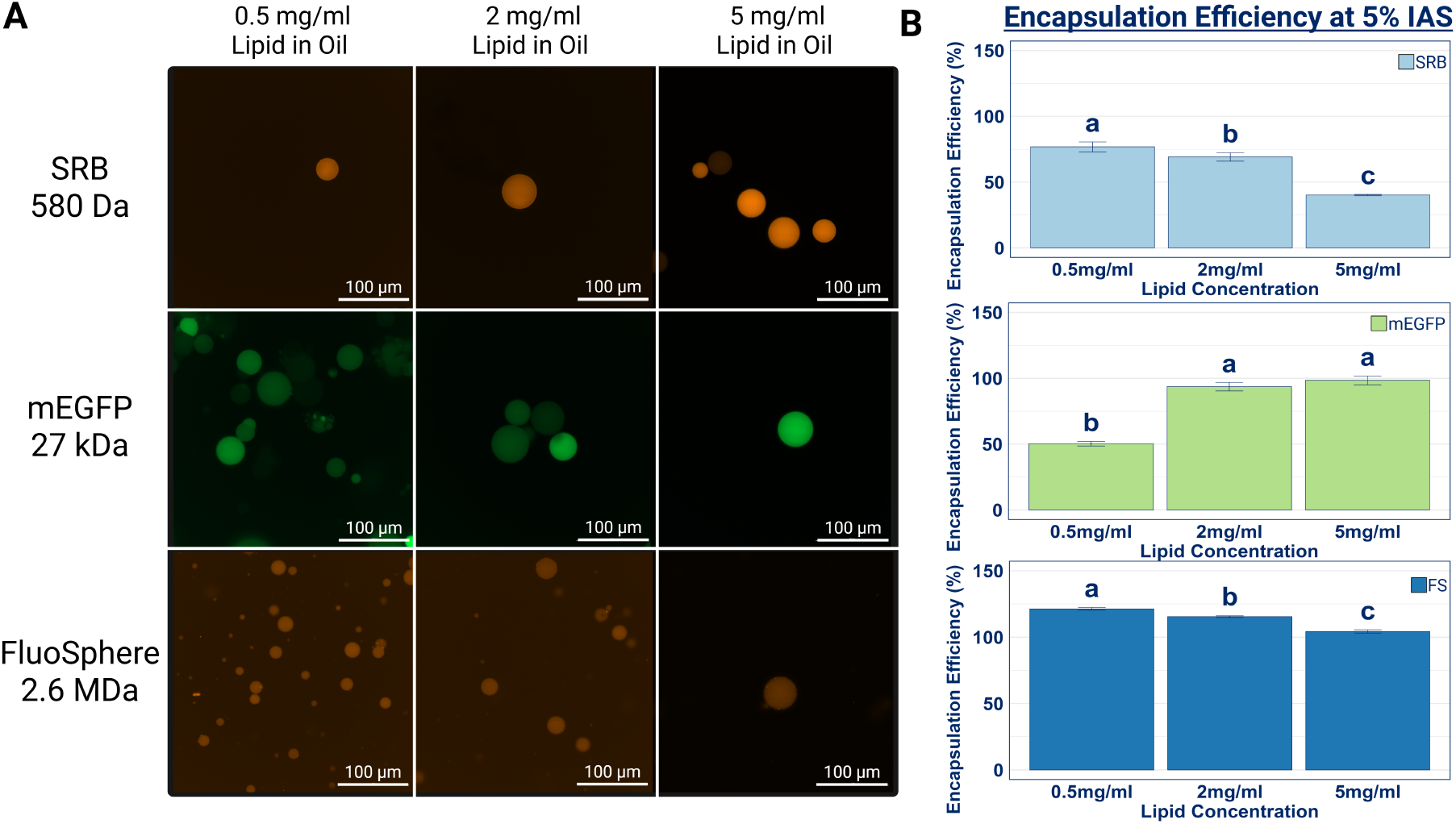
Encapsulation efficiency of fluorescent molecules of differing sizes (A) Representative fluorescence microscopy images of GUVs formed with varying lipid-in-oil concentrations (0.5, 2, and 5 mg/mL). Three molecules with different sizes -Sulforhodamine B (SRB, 580 Da), monomeric enhanced green fluorescent protein (mEGFP (27 kDa), and FluoSpheres (FS, 2.6 M Da) are encapsulated. (B) Quantification of encapsulation efficiencies with QuantGUV. The efficiency for SRB and FS significantly decreased with increasing lipid concentration while the encapsulation efficiency for mEGFP increased. Data are presented as mean ± standard error of the mean (SEM) from 3-5 biological replicates. Letters (a, b, c) denote statistically significant differences between groups as determined by a one-way ANOVA with a Tukey’s post-hoc test (p < 0.05) (See Supplementary Table S11 for full F-statistics and post-hoc p-values) Columns not sharing a letter are statistically significant to each other.

As expected, higher encapsulation efficiency was observed with increased size of molecules, with efficiency values ranging from 104.31 % to 121.36 % for FS (Figure 3B). Likely due to bigger molecules finding it difficult to diffuse out of the vesicles during the formation due to the slow diffusion and the difficulty in escaping through small defects such as spontaneous pores in the membrane. For mEGFP the encapsulation efficiency increased along increasing LiO concentrations. Specifically, efficiencies increased from 43.7 % at 0.5 *mg/mL* LiO to 88.4 % at 5 *mg/mL* LiO. This significant increase suggests that for the medium protein size range, higher LiO concentrations are favourable. In contrast to the mEGFP, SRB and FS showed a decreasing trend in encapsulation efficiency as the LiO concentration increased. The most significant change for SRB was observed changing between the highest and lowest concentrations, 0.5 *mg/mL* LiO to 5 *mg/mL* LiO, with a reduction of the mean encapsulation from 74.2 % to 38 %. This suggests that for smaller molecules, higher LiO concentrations lead to less efficient encapsulation. For FS, the encapsulation efficiency remained consistently high, with a mean encapsulation of 119.9 % for 0.5 *mg/mL* LiO which decreased to 103.2 % for 5 *mg/mL* LiO. The observed efficiencies greater than100 % may indicate aggregation of the styrene particles or a concentration effect within the vesicles. Although the overall decrease was not as statistically significant as SRB, the most significant change occurred between 0.5 *mg/mL* and 2 *mg/mL* LiO rather than between the highest and lowest observed for SRB. Despite the size difference this indicates a correlation between the small molecules and the larger molecules favouring lower lipid concentrations for enhanced encapsulation efficiency.

### 3.4 Effect of IAS Ratio and Lipid Concentration on Encapsulation Efficiency

To further gain insight on the encapsulation process, we changed the IAS ratio whilst maintaining the same LiO parameters and measured encapsulation efficiency of different molecules (shown in Supplementary information S8). Figure 4 presents the bar charts of the mean average encapsulation efficiency of each combination of IAS ratio and LiO concentration for each molecule. The results in figure 4 show the strong dependence of encapsulation efficiency on both the size of molecules and the concentration of lipids in the membrane, as evidenced by the large difference in efficiency for SRB (ranging from 35.2 % at 5 *mg/mL* LiO to 124.5 % at 0.5 *mg/mL* LiO). In contrast, a reduced dependence on the IAS ratio is shown throughout, particularly for FS, where the efficiency varied by only 2.8 percentage points 116.7 % to 119.5 %) at the optimal 0.5 *mg/mL* LiO concentration. mEGFP encapsulation efficiency continued to show the positive correlation with LiO concentration across all IAS ratios tested.The efficiency peaked at 5 *mg/mL* LiO across all IAS ratios. Despite the opposing correlations, similarly SRB encapsulation efficiency increased as IAS ratios were increased. The optimal encapsulation efficiency was observed at 5 *mg/mL* LiO, by increasing the IAS ratio from 1 % to 10 % an increase of encapsulation efficiency from 73.4 % to 91.8 % was observed. For the smaller molecule, SRB, the data exhibits two clear trends. Firstly, the same inverse relationship between LiO concentration and encapsulation efficiency is present for all IAS ratios tested, with the highest efficiencies consistently found at 0.5 *mg/mL* LiO. Secondly, the efficiency shows a positive correlation with the IAS ratio, this effect was more pronounced at the lower LiO concentrations with the greatest increase by increasing the IAS ratio from 1 % to 10 % the efficiency increased from 71.1 % to 124.5 % for 0.5 *mg/mL* LiO. The largest molecule, FS, exhibited a trend similar to SRB, with encapsulation efficiency anticorrelating with LiO (figure 4c). The highest encapsulation efficiency FS were consistently observed at 0.5 *mg/mL* LiO. However, unlike the smaller molecules, the influence of IAS ratio on FS encapsulation was less pronounced, with relatively high efficiencies maintained across all three IAS conditions. This further suggests that due to the size of the FS, leakage and other mechanisms of loss aren’t as prominent for larger species.

**Figure 4:**
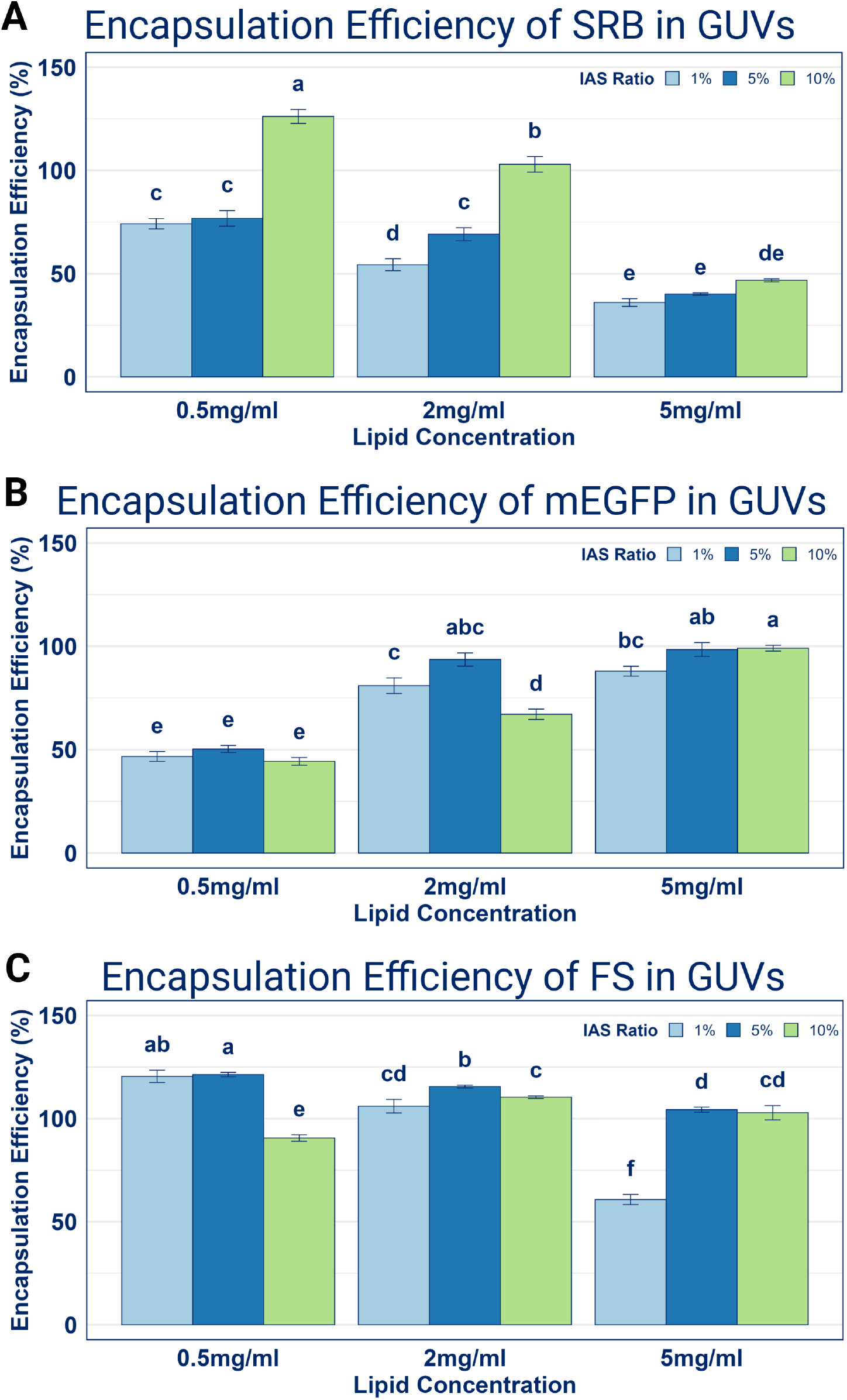
Encapsulation Efficiencies of varying IAS ratio in varied LiO concentrations The encapsulation efficiency of three different fluorescent molecules was assessed: (A) Sulforhodamine B (SRB), (B) monomeric Enhanced Green Fluorescent Protein (mEGFP), and (C) FluoSpheres (FS). GUVs were prepared using varying total lipid concentrations (0.5, 2, and 5 mg/mL) and different Inner Aqueous Solution (IAS) to lipid solution volume ratios (1, 5 and 10 %). Bars represent the mean encapsulation efficiency (%) from three to five independent experiments (n = 3 − 5), with error bars indicating ± standard error of the mean (SEM). Letters (a, b, c, d, e and f) denote statistically significant differences between groups as determined by A two-way ANOVA followed by Tukey’s post-hoc test was used for statistical analysis. Bars not sharing a common letter denote a statistically significant difference between the11groups (p < 0.05)(See Supplementary Table S12 for full F-statistics and post-hoc p-values).

### 3.5 Effect of Temperature and Molecular Crowding for Encapsulation

Molecular crowding is known to enhance biochemical reactions. To test its effect, we added different concentrations of polyethylene glycol (PEG) 8000 to the IAS to determine its impact on the concentration of mEGFP encapsulated(Figure 5A). 2 *mg/mL* LiO with a IAS ratio of 2.5 % was used at four PEG concentrations. Hence, GUVs were produced in PEG free and 3, 4 *and* 5% w/v PEG conditions (figure 5a). To correct for any fluorescence change caused by the addition of PEG to the system [47], Bulk fluorescence of IAS containing 6 µM mEGFP in the presence of PEG was measured at varying PEG concentrations. QuantGUV analysis of the vesicles show a significant increase in mEGFP encapsulation efficiency as PEG increases (figure 5B). For instance we saw an increase of encapsulation efficiency from 58.73 % at 0 % PEG to 107.27 % at 5 % PEG. This demonstrates a positive effect of crowding agents on the encapsulation efficiency of mEGFP We also tested the effect of maintaining low temperature during the GUV formation to the encapsulation efficiency during GUV formation. This is to maintain the activity of biochemical reactions during the encapsulation process. Two temperatures, 4 and 20^*°*^*C* were chosen to represent the shift from room temperature to a reduced temperature (Figure 5C). To ensure the fluorescence signal was not affected by the temperature during formation, images were taken at room temperature. QuantGUV analysis of the vesicles shows, the mean encapsulation efficiency decreased from 52.6 % at 20^*°*^*C* to 40.8 % at 4^*°*^*C* (Figure 5C, D). Whilst statistically significant, this decrease may be considered an acceptable compromise to preserve the functional integrity of encapsulated systems.

**Figure 5:**
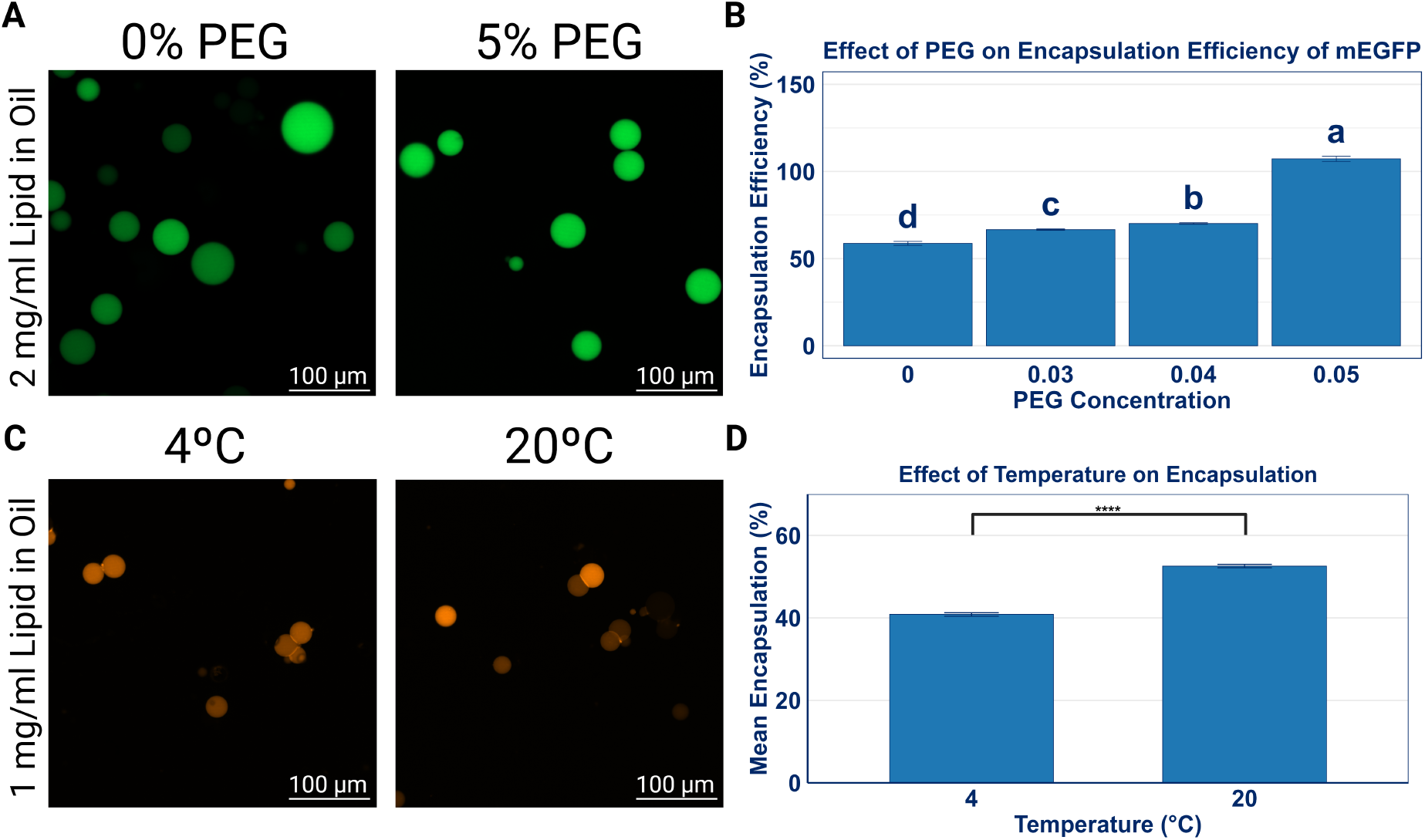
Effect of Temperature and PEG on GUV Encapsulation Efficiency The encapsulation efficiency of GUVs was explored further (A) Confocal images of GUVs containing in the presence of 0 % PEG, left, and 5 % PEG, right, Scale bars = 100 µm (B) Mean encapsulation efficiency of mEGFP encapsulated within GUVs using a variation of PEG concentrations 0, 3, 4 and 5% w/v using a 2 mg/mL LiO and a IAS ratio of 2.5 %. Data presented as mean ± standard error of the mean (SEM) from 3-5 biological repeats. Letters (a, b, c) denote statistically significant differences between groups as determined by a one-way ANOVA with a Tukey’s post-hoc test (p < 0.05) (See Supplementary Table S13 for full F-statistics and post-hoc p-values) (C) Sulforhodamine B (SRB), was encapsulated within GUVs using a formation temperature of 4, left, and 20^°^C, right, using a 1 mg/mL LiO and a IAS ratio of 2.5 %. (D) Mean Encapsulation of SRB presented as mean ± standard error of the mean (SEM) from 3-5 biological repeats, a two tail independent t-test was used to analyse significance, **** indicates p < 0.0001.(See Supplementary Table S14 for full statistics and p-values)

## 4 Method

### 4.1 Lipid in Oil Preparation

1,2-dioleoyl-sn-glycero-3-phosphoethanolamine-N-(biotinyl) (sodium salt) (Cat#870282) (18:1 Biotinyl DOPE) and 1,2-dioleoyl-sn-glycero-3-phosphocholine (Cat#850375) (18:1 DOPC) were purchased from Avanti Polar Lipids. Lipid dye 1,1’-Dioctadecyl-3,3,3’,3’-Tetramethylindodicarbocyanine, 4-Chlorobenzenesulfonate Salt (Cat#D7757) (DiD) was purchased from Thermo Fisher Scientific. Stock solutions in chloroform and ethanol were stored at *−*20^*°*^*C* Lipid in Oil (LiO) solutions were prepared by mixing 18:1 DOPC lipids (25 mg*/*mL), with 18:1 Biotinylated-DOPE lipids (1 mg*/*mL) and DiD (1 mg*/*mL) in 20 mL WHEATON® liquid scintillation vials (Cat#DWK986546) from Sigma Aldrich to a ratio of 99.8 : 0.1 : 0.1 % *w/v*. The solvent was desiccated in a fume hood under a dry stream of nitrogen whilst rotating to form a thin dry lipid film. The lipid film was then dispersed in 2 mL BioReagent mineral oil (Cat#M5904) obtained from Sigma Aldrich to achieve a final concentration of 0.5, 2, or 5 mg*/*mL. Glass vials were sealed with Parafilm, then LiO solutions were sonicated in a sonic bath at 40^*°*^*C* for 30 mins and were used the same day in experiments.

### 4.2 Inverted Emulsion Phase Transfer

The internal aqueous solution (IAS) containing 1x Phosphate Buffered Saline (PBS, Cat#10010023, Thermo Fisher Scientific) and 200 mM sucrose (Cat#A15583.36, Thermo Fisher Scientific) was added to the LiO solutions, the amount of IAS added in a ratio to the oil forms the IAS ratio, IAS ratios of 0.5, 2, or 5% were used. Typically 300 µL of LiO solution was used with a cosponsoring amount of IAS. The mix was then mechanically emulsified on a standard eppendorf rack at approximately 1 pass per second for 10 passes. The emulsion was then incubated at 4^*°*^*C* for 2 minutes. The external aqueous solution (EAS). 100 µL containing 1x PBS and 200 mM glucose (Cat#15023021, Thermo Fisher Scientific) was placed in a 1.5 mL eppendorf tube, and 150 µL of the emulsion was layered on top of the EAS. The stack was then centrifuged at 7000 g for 8 minutes. The transferred GUVs form a pellet which is collected by aspiration and redissolved in fresh EAS. For experiments using fluorescence dyes, the IAS/EAS is supplemented with 50 µM Sulforhodamine B (SRB, Cat#S1402, Sigma Aldrich), 6 mM monomeric enhanced green fluorescent protein (mEGFP, which was prepared as described in the Supplementary Information S15) or 0.5% *w/v* 20 *nm* FluoSpheres Carboxylate-Modified Microspheres (Cat#F8784, Thermo Fisher Scientific). For experiments to investigate the effect of varying temperature, Emulsions were prepared at 4 and 20^*°*^*C* using a 1 *mg/mL* LiO with a IAS ratio of 2.5 %. For the effect of crowding agents the IAS used for formation was supplemented with 3, 4 *and* 5% w/v PEG 8000 (Cat#043443.A3, Thermo Fisher Scientific) and sucrose concentrations lowered to maintain osmolarity. GUVs were made with a 2 *mg/mL* LiO with a IAS ratio of 2.5 %.

### 4.3 Microplate Surface Preparation

96 well microplate wells (Cat#165305, Thermo Fisher Scientific) were passivated to avoid adhesion and bursting of the GUVs. Wells were pre coated in 200 µL 5 mg*/*mL Bovine Serum Albumin (BSA, Cat#A9647, Sigma Aldrich) in 1x PBS containing 0.1 % Biotin labelled bovine albumin (Bio-BSA, Cat#A8549, Sigma Aldrich) by incubating for 30 minutes at 4^*°*^*C*. Wells were washed by removing 150 µL and replacing with 150 µL of 1x PBS. This was repeated a total of 4 times before 150 µL of 10 *ng/µL* NeutrAvidin (Cat#31055, Thermo fisher Scientific) was added and incubated for 30 minutes at 4^*°*^*C*. 150 µL was removed and washed with 150 µL of EAS for a total of 5 washes. Plates were stored at 4^*°*^*C* and were used the same day in experiments.

### 4.4 Confocal microscopy

Produced GUVs were collected by aspiration of oil layer and resuspension of GUV pellet in a treated 96 well plate. GUVs were imaged using a BioTek Cytation C10 Confocal Imaging Reader from Agilent Technologies. Images were acquired at 20x and 40x magnification using a 60 µm spinning disk confocal. The GFP, RFP, and Cy5 channels were utilized to detect fluorescent signals from membrane dyes (shown in Supplementary information S6) and encapsulated fluorophores. To allow for the direct comparison of fluorescence intensity, all images were acquired using a fixed gain of 3 and pixel intensities were normalised using the integration time for each image to a standard of 25 ms. To correct for nonlinear detectors a correction factor was applied (shown in Supplementary information S1), Flat field corrections were also applied to images to remove unequal illumination (shown in Supplementary information S2) Both TIF and PNG images were saved for processing and analysis. For each experimental condition, independent experiments were repeated between 3-5 times, GUVs were imaged and analysed using QuantGUV.

### 4.5 QuantGUV Image Analysis

Image analysis was performed using a custom graphical user interface (GUI) developed in Python, QuantGUV, available on Zenodo (10.5281/zenodo.17264460) and future updates available on Github (https://github.com/Marshall-Zak/QuantGUV). The program was designed to automate the high throughput detection and intensity quantification of GUVs from confocal TIF images.

Our analysis pipeline consisted of several steps. Firstly, images were pre-processed. Each image was converted to an 8 bit grayscale format, inverted, and then thresholded using Otsu’s thresholding method (cv2.THRESH OTSU). Potential GUVs were identified in the resulting binary image using a blob detection algorithm implemented with (cv2.SimpleBlobDetector create(params)). The blob detection algorithm isolates groups of bright pixels on a dark background and was used to detect potential GUVs containing fluorescent dyes. Detected blobs were then filtered based on defined parameters for area, circularity, and convexity. Detected objects were filtered to include only those with a size between 5 µm and 100 µm, a convexity greater than 0.9 and a circularity greater than 0.7. QuantGUV was first validated on user generated images (shown in Supplementary information S3). A circularity greater than 0.7 was applied to exclude objects with irregular or blurred boundaries. For each GUV that fits the parameters specified a 10 x 10 square pixel region of interest (ROI) was defined at the vesicles geometric centre, this ensures the mean measurement is representative of the vesicle lumen and minimises signal from potential membrane interactions. The detected GUVs and the defined ROIs were then quantified from the original unchanged TIF image containing the original observed dynamic range of pixel intensity.

Background correction was applied on an image by image basis. A binary mask of all detected GUVs in an image was created and inverted and assigned as the background area (shown in Supplementary information S4). The mean intensity of the lowest valued 10 % of pixels in this background region was calculated and subtracted from each GUV’s measured intensity. This method corrects for background noise and out of focus fluorescence while avoiding bias from bright aggregates and undetected, out of focus vesicles. Following automated analysis, an interactive review step within the GUI allowed for the manual inspection and exclusion of any remaining false positive detections before final data export. The final background corrected intensity for each GUV was then interpolated on a standard curve to estimate internal concentration.

### 4.6 QuantGUV: Standard Curve production

Bulk solutions of target fluorescent molecules were prepared in IAS and EAS. For bulk dye, 100 µL of IAS containing a concentration range of the fluorophores was used. Samples were placed into a 96 well microplate well and imaged on a Citation C10 confocal plater reader with Gain and integration time were fixed to 3 and 25 ms. Sample images were collected from multiple points in the well at a z height relevant to where the GUVs were observed, for our standard curve, the z value was set to 1900 µm. The mean average of all images in each fluorophores data set was used as the *I*_*Bulk*_ value for a given concentration.

For blank GUVs, GUVs were produced as described above in the absence of dye, GUVs were collected as before and resuspended in 100 µL of EAS containing a concentration range of fluorophores. Samples were placed into a 96 well microplate well and imaged on a Citation C10 confocal plater reader. Gain and integration time were again fixed to 3 and 25 ms. Images of the blank GUVs were analysed via QuantGUV using inverted TIFs to detect the blank GUVs as they appear as dark spots on a bright background. The same pipeline was used to detect and quantify intensity within the blank GUVs, however (cv2.equalizeHist(image)) and (cv2.GaussianBlur(equalized image, (11, 11), 0)) were used to enhance the contrast of images further to enable detection at low fluorophore concentrations. the internal signal of the blank GUVs was assigned to the *I*_*Blank*_ value and the background signal was assigned to the *I*_*Bulk*_ for the purpose of determining *I*_*Inside*_ for each GUV.

The calculated *I*_*Inside*_ values were plotted with the relevant concentration of fluorescent molecule to create standard curves (shown in Supplementary information S5). for each molecule blank IAS and EAS were used to sample a zero fluorescence condition to which the intercept of each standard curve was set.

### 4.7 Statistical Analysis

Statistical analysis was conducted using R Studio Build 513. To identify significant differences in GUV encapsulated concentrations at different LiO Concentrations (figure 3B), a one way ANOVA was carried out in R studio followed by a Tukey’s post-hoc test (*p <* 0.05) to identify groups of significant difference. For identifying significant differences in GUV encapsulated concentrations at different LiO Concentrations as IAS ratios (figure 4), a two way ANOVA was carried out in R studio followed by a Tukey’s post-hoc test (*p <* 0.05) to identify groups of significant difference. Significance was represented in figures 3B and 4 with the convention of letters (a, b, c, d, e and f) where columns that share one letter the same are not significantly different from each other whereas if no common letters are shared a significant difference was observed.

For identifying significance in the effect of PEG concentration on encapsulation efficiency (figure 5A), a one way ANOVA was carried out in R studio followed by a Tukey’s post-hoc test (*p <* 0.05) to identify groups of significant difference. Significance was represented in figure 5A with the convention of letters (a, b and c) where columns that share one letter the same are not significantly different from each other whereas if no common letters are shared a significant difference was observed.

For identifying significance in the effect of temperature on encapsulation efficiency, a two tailed independent t-test (Welch’s t-test) was conducted, threshold alpha was set at 0.05. Significance was shown with significance brackets with P values being represented in figure 5B using the convention of * indicates *p <* 0.05, ** indicates *p <* 0.01, *** indicates *p <* 0.001 and **** indicates *p <* 0.0001.

## 5 Discussion

In this work, we developed QuantGUV, a standardised, accessible software pipeline to quantify concentrations of fluorescent molecules encapsulated in GUVs at the single GUV level. By applying this high throughput method, we systematically investigated how various factors including physical properties of molecules, lipid concentration, IAS ratio, temperature and addition of crowding agents influence the encapsulation efficiency during the GUV formation. Our findings reveal that encapsulation of molecules inside of GUVs is not governed by a single parameter but is a complex interplay of factors, with molecular size being a particularly critical determining factor.

### 5.1 Molecular Size and Lipid Concentration Dictate Encapsulation Success

The size of the molecules has the biggest impact on the encapsulation efficiency with higher yield for bigger molecules. This is expected as 1. small defects in the lipid membrane that could form during the transfer or centrifugation process would not be sufficiently large to allow bigger molecules to leak and 2. 20 nm particles will diffuse 20 times slower than molecules with 1 nm in size. Also, there are indirect evidences that bigger molecules having higher encapsulation efficiencies as where addition of small molecules outside of GUVs were sufficient to achieve high yields for biochemical reactions [39]. Our work is the first attempt to provide direct experimental evidence on the relationship between the size of molecules and their encapsulation efficiencies.

We note that opposing trends in encapsulation efficiency were observed for different sized molecules when lipid concentration in the oil has changed. mEGFP showed higher encapsulation efficiencies for higher lipid concentrations, while SRB and fluospheres showed the opposite trends with the effect more prominent for SRB. In general, higher lipid concentration would prevent defects on the lipid membranes and faster healing when defects occur, potentially leading to higher encapsulation efficiencies. The encapsulation of mEGFP fits to this trend while the other two molecules do not. As the encapsulation efficiencies of SRB increased when we increase the IAS ratio, which could further destabilise the lipid membrane by making lipid more sparse, we speculated that the interaction with lipid membrane could affect the encapsulation efficiencies. When we examined the images of GUVs encapsulating SRB and fluospheres, we observed a distinct halo of fluorescence at the GUV membrane (shown in Supplementary information S7), indicating the sequestration of SRB at the lipid membrane. No halo was observed for GUVs encapsulating mEGFP. Therefore, depending on the properties of the molecules, the membrane interface could act as a sink, depleting the molecules in the GUVs lumen. mEGFP, being a highly polar protein, does not exhibit this membrane sequestration. Its likley that the monomeric A206K mutation, which replaces a hydrophobic residue with a charged lysine, further ensures the protein remains aqueous by eliminating a potential hydrophobic interaction patch, thereby preventing it from partitioning to the lipid interface.

### 5.2 The Influence of IAS Ratio on Encapsulation Efficiency

Our results show that encapsulation efficiency of SRB improves with an increasing IAS ratio from 1 % to 10 %. However, mixed results were observed for other molecules and therefore we cannot attribute this behaviour to the effect of the size of molecules. However, increasing the IAS ratio could increase the number of vesicles produced and therby transferring more molecules to the lumen of GUVs.

### 5.3 Effect of varied temperature and crowding agents on encapsulation

Reducing the temperature had a minor negative impact on the encapsulation efficiency of mEGFP while addition of PEG increased the encapsulation efficiency significantly. Forming synthetic cells at reduced temperatures presents a strategic trade off in our results, accepting a modest decrease in encapsulation efficiency in order to stall the encapsulated biological reactions. Given that the sample preparation and mixing process takes approximately 30 mins which is long enough for many reactions to initiate. A low temperature approach at 4^*°*^*C* is preferred for precise implementation of most synthetic systems. A plausible physical mechanism for this lower encapsulation efficiency stems from the temperature dependent properties of the oil phase. As temperature decreases, the viscosity of the LiO mixture and the lipid interface increases, which could induce more significant deformation and slower recovery of defects. The positive impact of the addition of PEG could come from the exclusion volume effect which would entropically drive mEGFP into the lumen of droplets. The diffusion of mEGFP would also slow down in the presence of crowding agent and the PEG could interact with the mEGFP to help with the encapsulation efficiencies. As the presence of PEG also increases the rates of biochemical reactions [39], it stresses the benefit of including molecular crowding agents in synthetic cell systems.

### 5.4 Limitations and Future Outlook

While the QuantGUV is a powerful tool, but it’s important to acknowledge its limitations coming from using fluorescence intensity as a main measurement. The fluorescence signal depends not only on the setup, it also depends on the local environment with pH, buffer composition, protein folding and aggregation affecting quantum yield of a fluorescent molecule. The standard curve calibration, while robust, assumes that light scattering by blank GUVs doesn’t significantly alter the out of focus light profile compared to a pure bulk solution. However, one thing to note is that our calibration process would work with most of the fluorescence microscopy techniques as in principle, the standard curve takes account of all the effects we described. Therefore, it should be possible to use our procedure in a standard wide-field microscope, although the effect of background would be significant. Therefore, by being a low barrier to entry tool, the QuantGUV has a potential to be applied in various systems to move beyond trial and error and open up building systematically engineering synthetic cells with predictable internal concentrations that allows for example, much more accurate mathematical modellings. Future work should focus on using this tool to explore a wider range of lipid compositions such as incorporating cholesterol or charged lipids and to co encapsulate multiple components in the same formation, paving the way for the reliable construction of complex, multipart biological systems within GUVs.

## Acknowledgements

The authors acknowledge funding from the Engineering and Physical Sciences Research Council (EPSRC) under grant number [EP/X023303/1], Biotechnology and Biological Sciences Research Council (BBSRC) under grant number [BB/X01262X/1] and Royal Society [RGS\R2\242552]. Z.M acknowledges the University of Surrey for funding the PhD studentship. The Authors also thank Dr. Agata Gajewicz-Jaromin for technical support

