## Supplementary information for "QuantGUV: Quantifying Encapsulation Efficiency of Small Molecules in GUVs"

### 1 Supplementary information

**Contents:**

| | |
|---|---|
| QuantGUV Calibration and Tests | S1-S5 |
| Merged Channels, DiD and each fluorophore | S6 |
| Halo effect at GUV membrane | S7 |
| GUV images at all conditions in figure 4 | S8 |
| Heterogeneity of encapsulation efficiency in figure 4 | S9 |
| GUV diameter distribution in figure 4 | S10 |
| Statistical Analysis Data | S11-S14 |
| GFP Production Protocol | S15 |

S1: integration time is an intensity per pixel per second factor, allowing for the normalization by the division of the integration time used and multiplied by 25 to achieve consistency with the standard curve. However detectors are non linear so a minor adjustment must be made to account for this effect

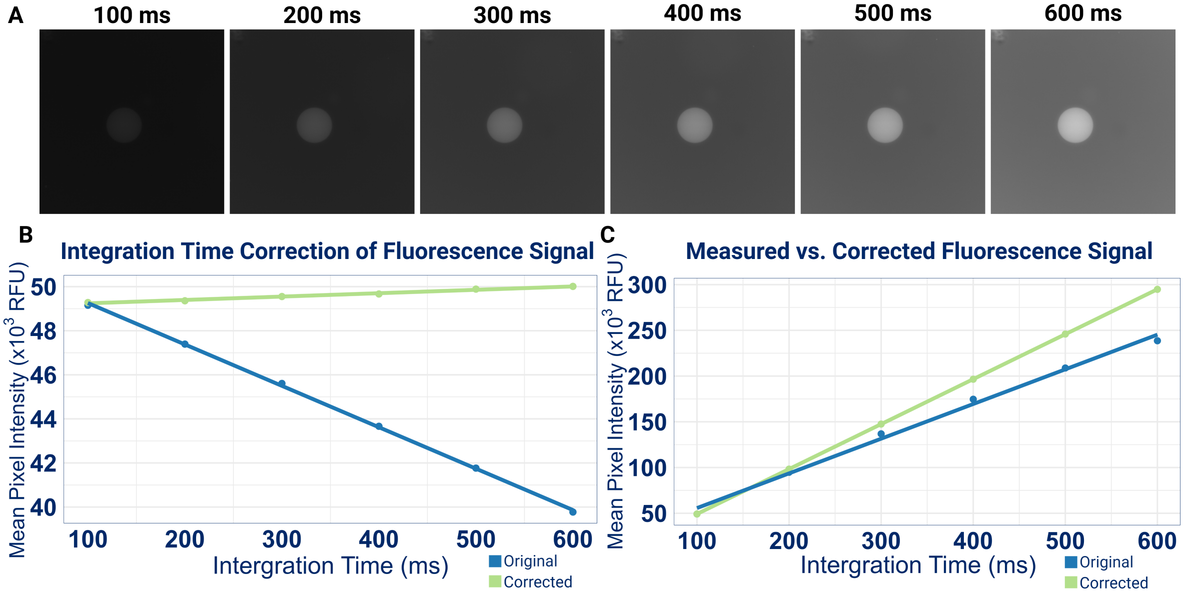


*S1: Integration time correction of fluorescence intensity A a time series of a GUV images as integration time increases B Pixel intensity was linearised through an empirical correction factor generated using integration times in a Polynomial Regression C Empirical correction at non normalised intensities showing linearisation of detected signal*

S2: Detected images were flattened using a flat fielding correction factor applied to all images before quantification.

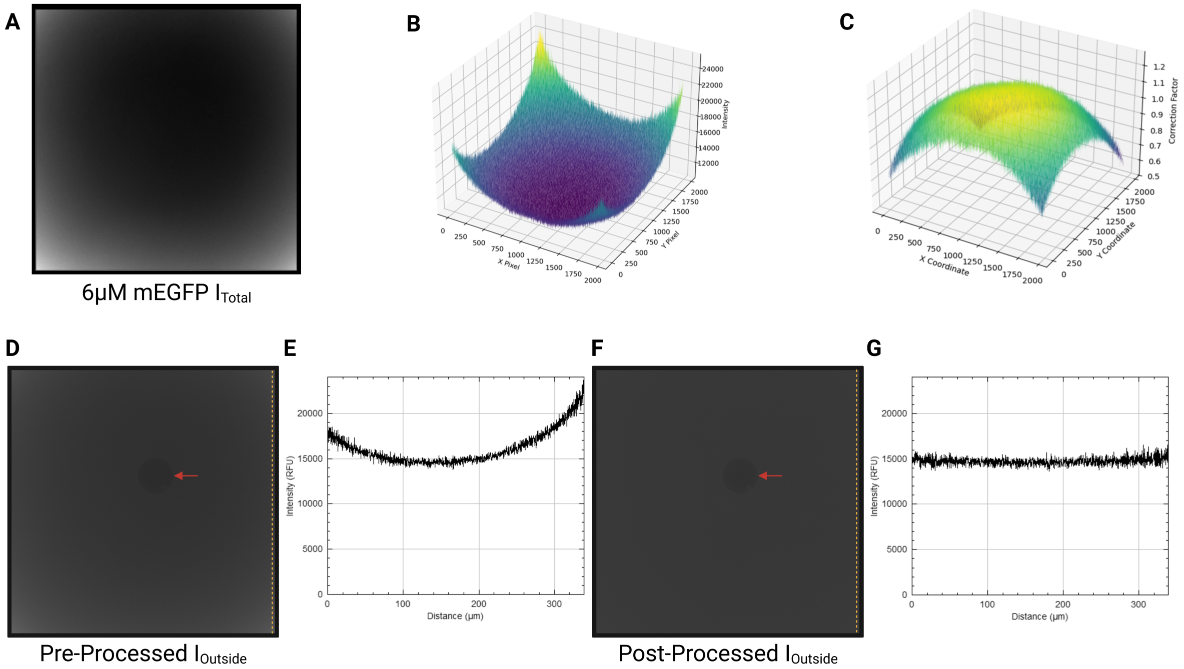


***S2: Detector normalisation through flat field processingA Bulk images of each fluorescent dye at max concentration were obtained, Tif placed on Black square to highlight vignette from detector B 3D pixel mapping to inspect vignette observed C 3D coordinate map of correction factors, produced from taking a mean pixel intensity from B and applying a factor to flatten the image D Pre-Processed $I_{Outside}$ image, Red arrow indicated a blank GUV, yellow dashed line indicates location of intensity profile plot E Intensity profile plot for D F Post-Processed $I_{Outside}$ image, Red arrow indicated a blank GUV, yellow dashed line indicates location of Intensity profile plot G Intensity profile plot for F***

S3: Test images were generated using python code to produce circles of varying size, circularity and intensity on a background intensity of users choice to test the calculations of QuantGUV on known parameters

A

```
# --- Image Settings ---
IMAGE_WIDTH = 1960
IMAGE_HEIGHT = 1960
BACKGROUND_INTENSITY = 5000
circles_properties = [
    # GUV1:
    {'radius_y': 28.9,  'radius_x': 28.9,  'intensity': 25000},
    # GUV2:
    {'radius_y': 57.8,  'radius_x': 57.8,  'intensity': 30000},
    # GUV3:
    {'radius_y': 86.7,  'radius_x': 86.7,  'intensity': 35000},
    # GUV4:
    {'radius_y': 115.6, 'radius_x': 115.6, 'intensity': 40000},
    # GUV5:
    {'radius_y': 144.5,  'radius_x': 144.5,  'intensity': 45000},
```

B

| GUV ID | Diameter (μm) | Intensity | Background Normalised |
|---|---|---|---|
| GUV 1 | 10 | 25000 | 20000 |
| GUV 2 | 20 | 30000 | 25000 |
| GUV 3 | 30 | 35000 | 30000 |
| GUV 4 | 40 | 40000 | 35000 |
| GUV 5 | 50 | 45000 | 40000 |

C D

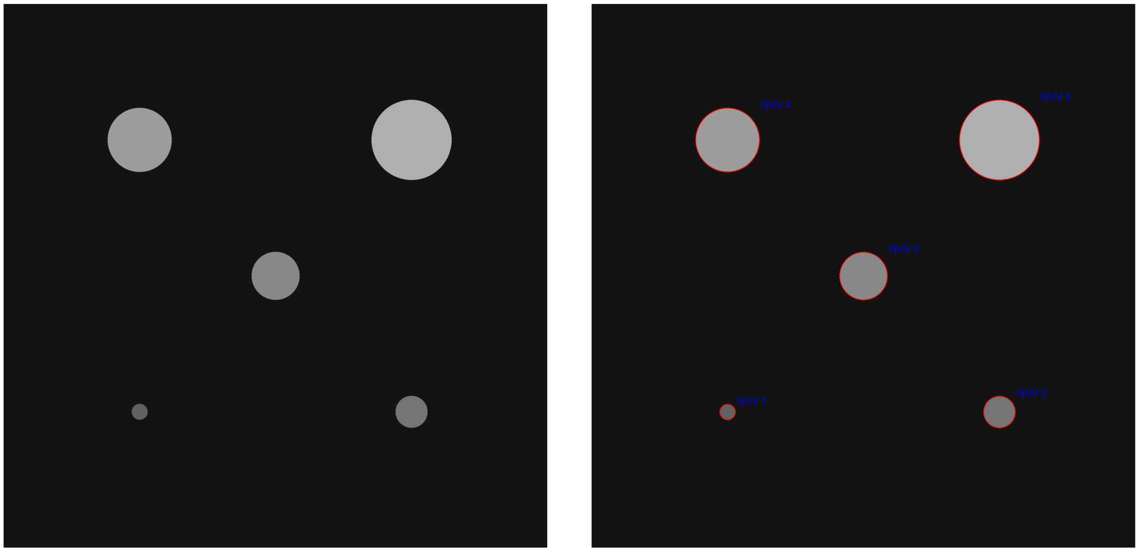

*S3: QuantGUV test image calibration A Custom code determining parameters of test images B Output from QuantGUV indicating correct calculations and subtractions were performed C Test TIF created an provided to QuantGUV D Output PNG from QuantGUv showing "GUV" detection*

S4: Image per Image background correction is built into QuantGUV, a mask of the detected vesicles that fit a parameter is produced and inverted to create a background mask, isolating all pixels associated with the background.

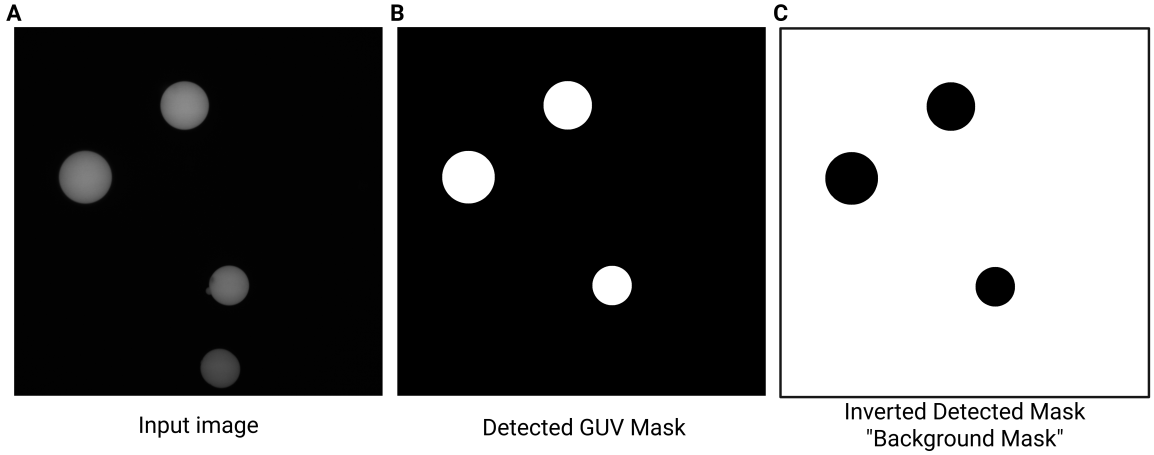


*S4: Background mask generationA Input TIF image provided to QuantGUV B Output PNG mask of detected GUVs C Output PNG of background mask, inverted from B*

S5: Standard curves were produced as described in main text, a standard curve was produced for each magnification to eliminate variation in intensity caused by magnification differences in data sets allowing for more insights at the individual level and population level for all conditions

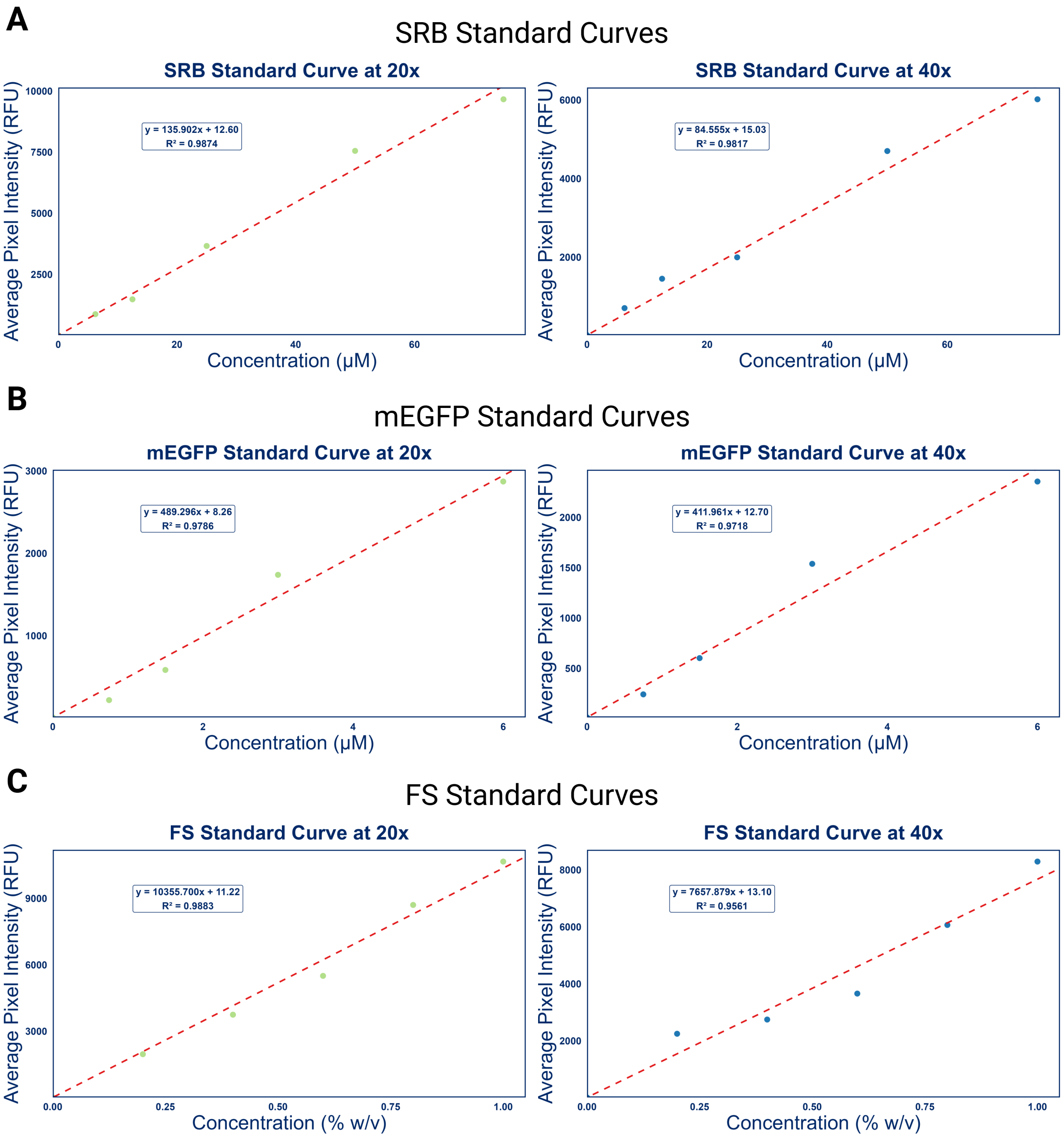


*S5: Standard Curves for fluorophores at each magnificationA SRB 20x Standard curve B SRB 40x Standard curve C mEGFP 20x Standard curve D mEGFP 40x Standard curve E FS 20x Standard curve F FS 40x Standard curve*

S6: DiD was added to all LiO preparations to allow for screening of GUVs, Merged Images were collected to show internal volume signal as well as membrane bound signal.

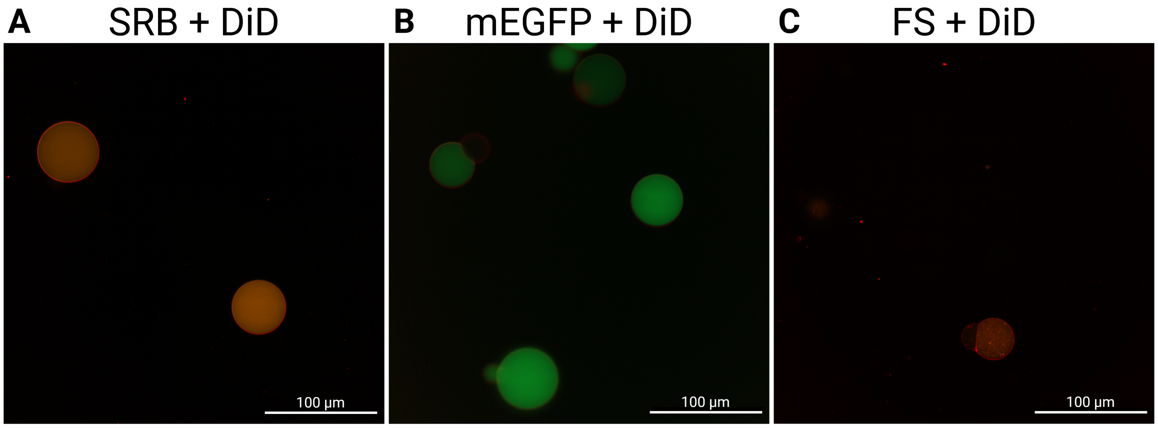


***S6: Merged GUV ImagesA Merged GUV image of DiD and SRB B Merged GUV image of DiD and mEGFP C Merged GUV image of DiD and FS. Scale bars*** = 100 μm

S7: A Halo Effect was observed in some conditions of SRB and FS, Intensity profile plots were used to inspect and visualise this effect at the membrane of GUVs

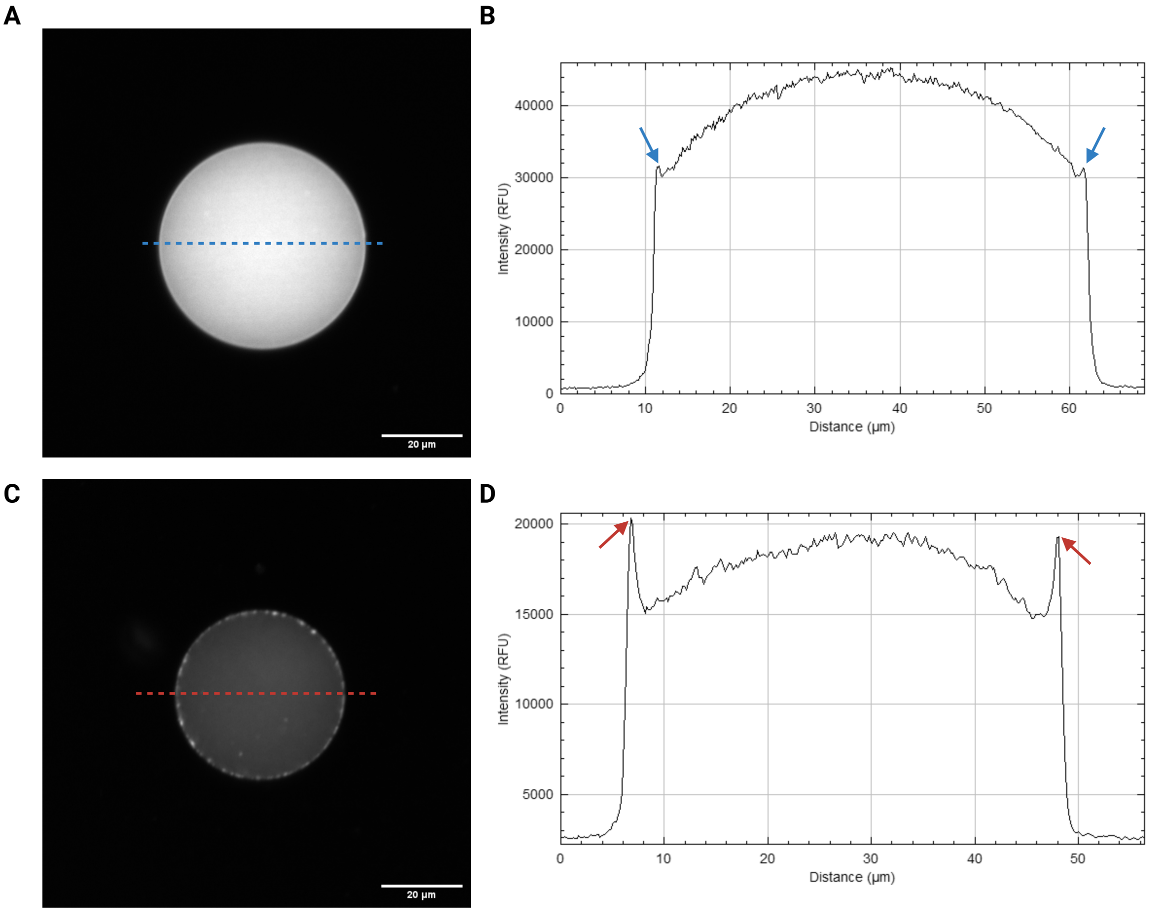


***S7: Halo Effect in GUVs containing SRB and FSA Confocal image of GUv containing SRB with a halo effect at the membrane, Blue line represents the location of Intensity profile plot B Intensity profile plot of A, Blue arrows indicate the Halo effect in the cross section of GUV C Confocal image of GUv containing FS with a halo effect at the membrane, Red line represents the location of Intensity profile plot D Intensity profile plot of C, Red arrows indicate the Halo effect in the cross section of GUV***

S8: Representative confocal images of each condition tested in figure 4, containing a matrix of LiO concentrations and IAS ratio per fluorescent molecules

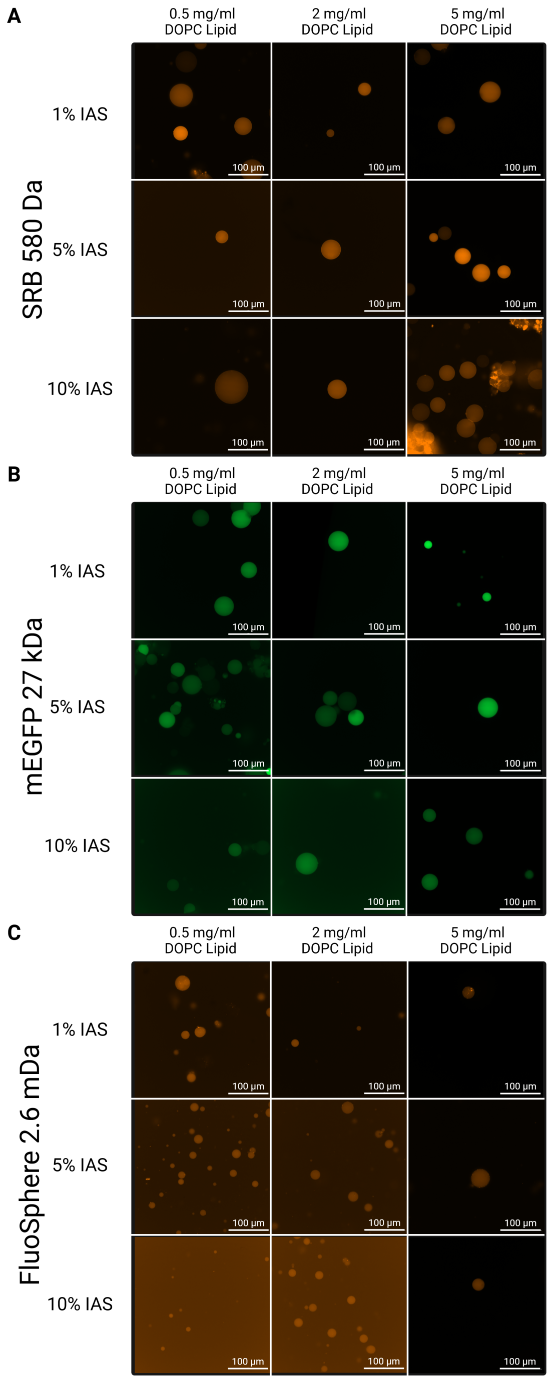


***S8: Confocal images of GUVs in conditions in figure 4A SRB confocal image matrix of GUVs containing SRB 580 Da at LiO concentrations of*** 0.5, 2 *and* 5 *mg/ml* ***in combination with IAS ratios of*** 1, 5 *and* 10% ***B mEGFP confocal image matrix of GUVs containing mEGFP 27 kDa at LiO concentrations of*** 0.5, 2 *and* 5 *mg/ml* ***in combination with IAS ratios of*** 1, 5 *and* 10% ***C FS confocal image matrix of GUVs containing FS 2.6 MDa at LiO concentrations of*** 0.5, 2 *and* 5 *mg/ml* ***in combination with IAS ratios of*** 1, 5 *and* 10%

S9: Box and whisker blots displaying the distribution of Encapsulation efficiency of each condition tested in figure 4, containing a matrix of LiO concentrations and IAS ratio per fluorescent molecules

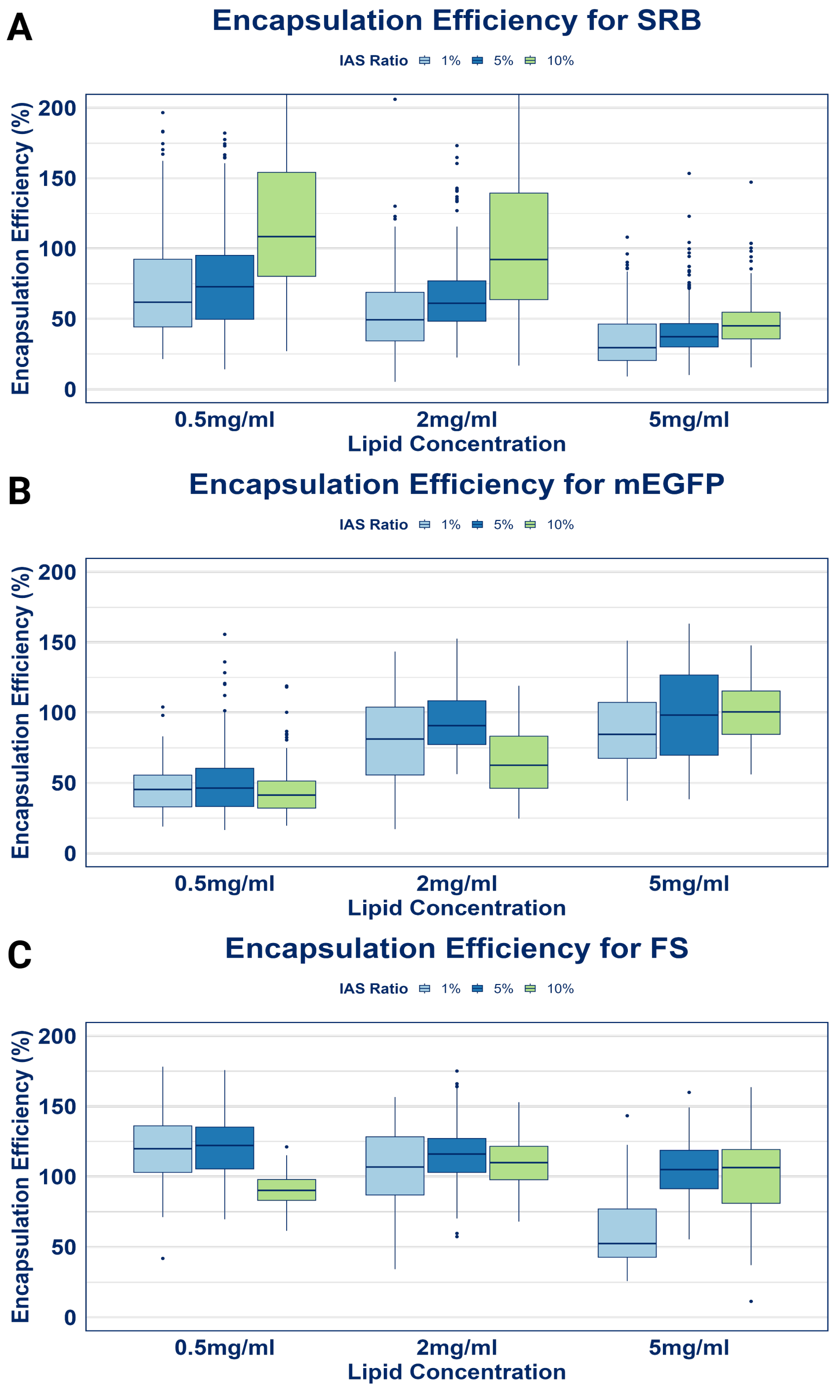


*S9: Encapsulation efficiency distributionsA Box and whisker plot of SRB encapsulation efficiencies of GUVs containing SRB 580 Da at LiO concentrations of* $0.5,\ 2\ and\ 5\ mg/ml$ *in combination with IAS ratios of* $1,\ 5\ and\ 10\%$ *B Box and whisker plot of mEGFP encapsulation efficiencies of GUVs containing mEGFP 27 kDa at LiO concentrations of* $0.5,\ 2\ and\ 5\ mg/ml$ *in combination with IAS ratios of* $1,\ 5\ and\ 10\%$ *C Box and whisker plot of FS encapsulation efficiencies of GUVs containing FS 2.6 MDa at LiO concentrations of* $0.5,\ 2\ and\ 5\ mg/ml$ *in combination with IAS ratios of* $1,\ 5\ and\ 10\%$

S10: Violin plot displaying GUV diameter distribution for each condition tested in figure 4, containing a matrix of LiO concentrations and IAS ratio per fluorescent molecules

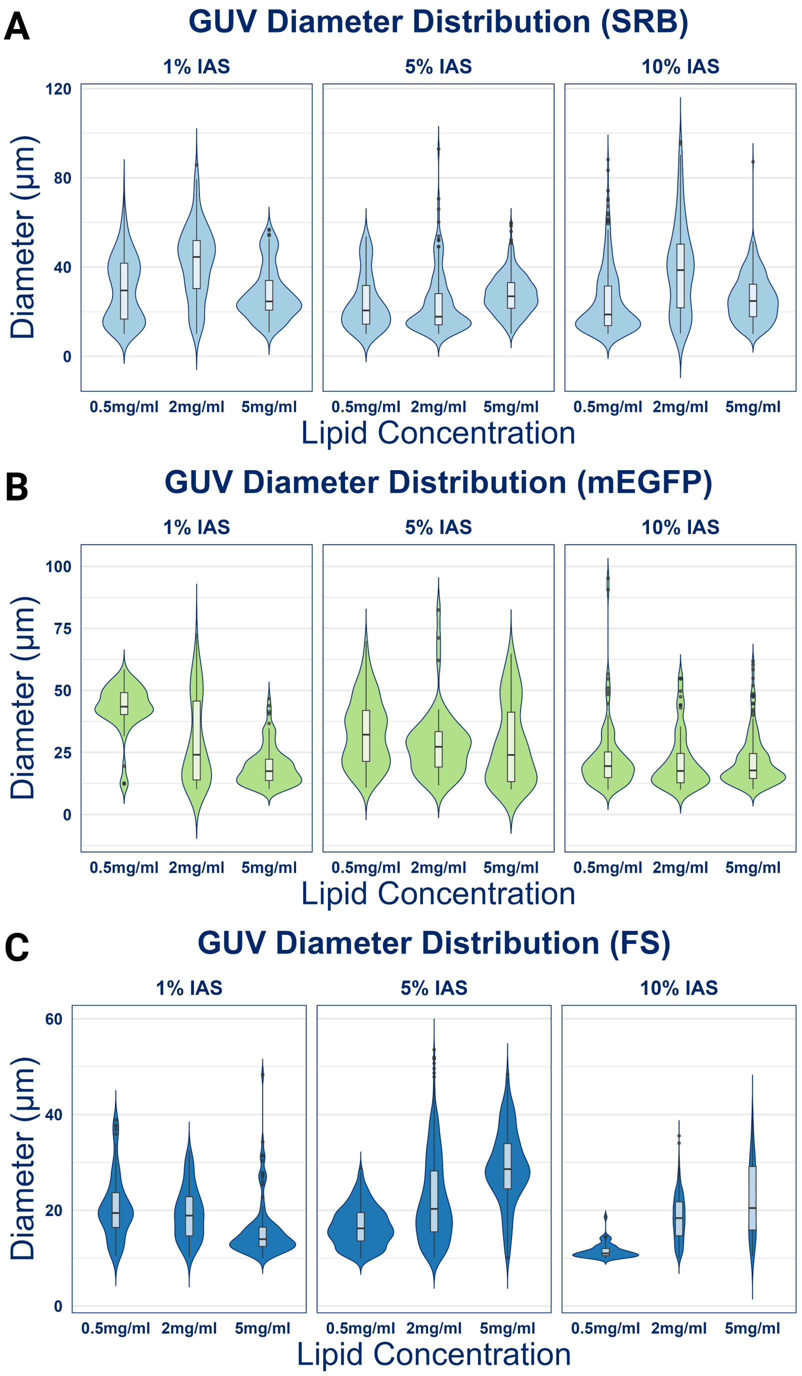


***S10: Violin plot of GUV diametersA Violin plot of GUV diameter for vesicles containing SRB 580 Da at LiO concentrations of*** $0.5,\ 2\ and\ 5\ mg/ml$ ***in combination with IAS ratios of*** $1,\ 5\ and\ 10\%$ ***B Violin plot of GUV diameter for vesicles containing mEGFP 27 kDa at LiO concentrations of*** $0.5,\ 2\ and\ 5\ mg/ml$ ***in combination with IAS ratios of*** $1,\ 5\ and\ 10\%$ ***C Violin plot of GUV diameter for vesicles containing FS 2.6 MDa at LiO concentrations of*** $0.5,\ 2\ and\ 5\ mg/ml$ ***in combination with IAS ratios of*** $1,\ 5\ and\ 10\%$

S11: Statistical Analysis results for data presented in figure 3. Containing statistical data from a one way ANOVA and post hoc test for each fluorophore

**A** Lipid effect on SRB Encapsulation
One Way ANOVA Summary

| Lipid Conc | N | Mean | SD | Significance Group |
|---|---|---|---|---|
| 0.5mg/ml | 121 | 76.81 | 41.33 | a |
| 2mg/ml | 156 | 69.13 | 39.76 | b |
| 5mg/ml | 675 | 40.18 | 14.87 | c |

**B** Lipid effect on SRB Encapsulation
PostHoc Summary

| term | df | sumsq | meansq | statistic | p.value |
|---|---|---|---|---|---|
| Lipid Conc | 2 | 208936.52 | 104468.26 | 165.50 | 2.19E-62 |
| Residuals | 949 | 599027.50 | 631.22 | NA | NA |

**C** Lipid effect on mEGFP Encapsulation
One Way ANOVA Summary

| Lipid Conc | N | Mean | SD | Significance Group |
|---|---|---|---|---|
| 0.5mg/ml | 90 | 98.45 | 31.86 | a |
| 2mg/ml | 52 | 93.63 | 23.15 | a |
| 5mg/ml | 185 | 50.34 | 23.61 | b |

**D** Lipid effect on mEGFP Encapsulation
PostHoc Summary

| term | df | sumsq | meansq | statistic | p.value |
|---|---|---|---|---|---|
| Lipid Conc | 2 | 173335.64 | 86667.82 | 127.48 | 1.44E-41 |
| Residuals | 324 | 220266.49 | 679.83 | NA | NA |

**E** Lipid effect on FS Encapsulation
One Way ANOVA Summary

| Lipid Conc | N | Mean | SD | Significance Group |
|---|---|---|---|---|
| 0.5mg/ml | 359 | 121.36 | 20.10 | a |
| 2mg/ml | 728 | 115.50 | 18.00 | b |
| 5mg/ml | 228 | 104.31 | 18.80 | c |

**F** Lipid effect on FS Encapsulation
PostHoc Summary

| term | df | sumsq | meansq | statistic | p.value |
|---|---|---|---|---|---|
| Lipid Conc | 2 | 40725.68 | 20362.84 | 58.04 | 7.01E-25 |
| Residuals | 1312 | 460283.54 | 350.83 | NA | NA |

*S11: Figure 3 Statistical AnalysisA Table containing statistic summary for a one way ANOVA comparing LiO effect on SRB encapsulation efficiency B Table containing statistic summary for a Tukey's post-hoc test comparing the One way ANOVA on LiO effect on SRB encapsulation efficiency C Table containing statistic summary for a one way ANOVA comparing LiO effect on mEGFP encapsulation efficiency D Table containing statistic summary for a Tukey's post-hoc test comparing the One way ANOVA on LiO effect on mEGFP encapsulation efficiency E Table containing statistic summary for a one way ANOVA comparing LiO effect on FS encapsulation efficiency F Table containing statistic summary for a Tukey's post-hoc test comparing the One way ANOVA on LiO effect on FS encapsulation efficiency*

S12: Statistical Analysis results for data presented in figure 4. Containing statistical data from a two way ANOVA and post hoc test for each fluorophore

**A** SRB 2 Way ANOVA Summary

| term | df | sumsq | meansq | statistic | p.value |
|---|---|---|---|---|---|
| Lipid Conc | 2 | 1623527.48 | 811763.74 | 620.66 | 1.49E-218 |
| IAS | 2 | 494298.16 | 247149.08 | 188.96 | 5.15E-77 |
| Lipid Conc:IAS | 4 | 191933.58 | 47983.40 | 36.69 | 8.01E-30 |
| Residuals | 2435 | 3184763.05 | 1307.91 | NA | NA |

**B** SRB PostHoc Summary

| Lipid Conc | IAS | N | Mean | SD | Significance Group |
|---|---|---|---|---|---|
| 0.5 mg/ml | 10% | 356 | 126.19 | 63.95 | a |
| 2 mg/ml | 10% | 169 | 103.01 | 49.44 | b |
| 0.5 mg/ml | 5% | 121 | 76.81 | 41.33 | c |
| 0.5 mg/ml | 1% | 250 | 74.19 | 40.13 | c |
| 2 mg/ml | 5% | 156 | 69.13 | 39.76 | c |
| 2 mg/ml | 1% | 135 | 54.36 | 33.45 | d |
| 5 mg/ml | 10% | 444 | 46.78 | 15.38 | de |
| 5 mg/ml | 5% | 675 | 40.18 | 14.87 | e |
| 5 mg/ml | 1% | 138 | 36.04 | 22.22 | e |

**C** mEGFP 2 Way ANOVA Summary

| term | df | sumsq | meansq | statistic | p.value |
|---|---|---|---|---|---|
| Lipid Conc | 2 | 452596.60 | 226298.30 | 381.82 | 5.09E-124 |
| IAS | 2 | 11586.03 | 5793.01 | 9.77 | 6.25E-05 |
| Lipid Conc:IAS | 4 | 26027.30 | 6506.83 | 10.98 | 1.02E-08 |
| Residuals | 1003 | 594454.19 | 592.68 | NA | NA |

**D** mEGFP PostHoc Summary

| Lipid Conc | IAS | N | Mean | SD | Significance Group |
|---|---|---|---|---|---|
| 5 mg/ml | 10% | 225 | 99.11 | 20.84 | a |
| 5 mg/ml | 5% | 90 | 98.45 | 31.86 | ab |
| 2 mg/ml | 5% | 52 | 93.63 | 23.15 | abc |
| 5 mg/ml | 1% | 119 | 88.00 | 25.87 | bc |
| 2 mg/ml | 1% | 76 | 80.98 | 32.84 | c |
| 2 mg/ml | 10% | 96 | 67.14 | 24.47 | d |
| 0.5 mg/ml | 5% | 185 | 50.34 | 23.61 | e |
| 0.5 mg/ml | 1% | 59 | 46.70 | 18.26 | e |
| 0.5 mg/ml | 10% | 110 | 44.35 | 19.35 | e |

**E** FS 2 Way ANOVA Summary

| term | df | sumsq | meansq | statistic | p.value |
|---|---|---|---|---|---|
| Lipid Conc | 2 | 150878.99 | 75439.49 | 178.51 | 1.48E-73 |
| IAS | 2 | 85121.42 | 42560.71 | 100.71 | 6.47E-43 |
| Lipid Conc:IAS | 4 | 111978.39 | 27994.60 | 66.24 | 1.55E-53 |
| Residuals | 2710 | 1145249.88 | 422.60 | NA | NA |

**F** FS PostHoc Summary

| Lipid Conc | IAS | N | Mean | SD | Significance Group |
|---|---|---|---|---|---|
| 0.5mg/ml | 5% | 359 | 121.36 | 20.10 | a |
| 0.5mg/ml | 1% | 73 | 120.49 | 25.51 | ab |
| 2mg/ml | 5% | 728 | 115.50 | 18.00 | b |
| 2mg/ml | 10% | 1025 | 110.38 | 20.65 | c |
| 2mg/ml | 1% | 84 | 106.02 | 30.20 | cd |
| 5mg/ml | 5% | 228 | 104.31 | 18.80 | d |
| 5mg/ml | 10% | 71 | 102.86 | 29.32 | cd |
| 0.5mg/ml | 10% | 52 | 90.57 | 11.13 | e |
| 5mg/ml | 1% | 99 | 60.76 | 24.63 | f |

*S11: Figure 4 Statistical AnalysisA Table containing statistic summary for a two way ANOVA comparing LiO effect on SRB encapsulation efficiency B Table containing statistic summary for a Tukey's post-hoc test comparing the two way ANOVA on LiO effect on SRB encapsulation efficiency C Table containing statistic summary for a two way ANOVA comparing LiO effect on mEGFP encapsulation efficiency D Table containing statistic summary for a Tukey's post-hoc test comparing the two way ANOVA on LiO effect on mEGFP encapsulation efficiency E Table containing statistic summary for a two way ANOVA comparing LiO effect on FS encapsulation efficiency F Table containing statistic summary for a Tukey's post-hoc test comparing the two way ANOVA on LiO effect on FS encapsulation efficiency*

S13: Statistical Analysis results for data presented in figure 5A. Containing statistical data from a One way ANOVA and post hoc test for the effect of PEG on mEGFP encapsulation efficiency

**A** PEG Concentration one Way ANOVA Summary

| term | df | sumsq | meansq | statistic | p.value |
|---|---|---|---|---|---|
| PEG_Conc | 3 | 318016.83 | 106005.61 | 373.45 | 3.88E-210 |
| Residuals | 3463 | 982981.61 | 283.85 | NA | NA |

**B** PEG Concentration PostHoc Summary

| PEG Conc | N | Mean | SD | Significance Group |
|---|---|---|---|---|
| 5% | 177 | 107.27 | 19.52 | a |
| 4% | 1182 | 70.18 | 14.89 | b |
| 3% | 1653 | 66.63 | 15.28 | c |
| 0% | 455 | 58.73 | 24.32 | d |

*S11: Figure 5A Statistical AnalysisA Table containing statistic summary for a one way ANOVA comparing the effect of PEG concentration mEGFP encapsulation efficiency B Table containing statistic summary for a Tukey's post-hoc test comparing the one way ANOVA on the effect of PEG concentration mEGFP encapsulation efficiency*

S14: Statistical Analysis results for data presented in figure 5B. Containing statistical data from a two tail independent T-Test for the effect of temperature on SRB encapsulation efficiency

| Variable Compared | Group 1 | Group 2 | t-statistic (t) | Degrees of Freedom (df) | Sample Size (n1,n2) | p-value (p) |
|---|---|---|---|---|---|---|
| Encapsulation Percent | 4 | 20 | -19.9 | 2825 | 1406, 1444 | < 0.001 |

*S11: Figure 5B Statistical Analysis A table containing statistical data from a two tail independant T-Test for the effect of temperature on SRB encapsulation efficiency*

S15: GFP Production Protocol pRSET his-eGFP was a gift from Jeanne Stachowiak Addgene plasmid # 113551 http://n2t.net/addgene:113551 ; RRID: Addgene_113551.

Plasmids were transformed into competent E. coli (BL21) cells and single colonies were selected to inoculate a 50 mL culture of LB (100 µg/mL ampicillin) at $37°C$, 250 RPM. Protein production was induced with the addition of IPTG to a final concentration of 1 mM and incubated for 16 h at $30°C$, 250 RPM. Bacteria were harvested by centrifugation at 4000 RPM and the pellets stored at $-80°C$.

The cell pellet was resuspended in B-Per complete bacterial protein extraction reagent (ThermoFisher) and centrifuged at 4000 RPM for 15 min. Purification was performed by HisPur superflow Ni-NTA agarose beads under conditions described by the manufacturer (thermoscientific) using Pierce™ Disposable Columns, 5 mL. Purified protein was eluted with 250 mM imidazole, the elution buffer was exchanged to PBS and concentrated with Amicon Ultra Centrifugal filters (Millipore Ultracel-10 K) with a 10 kDa cut off. Concentrated samples in PBS were stored at $4°C$.Samples were concentrated to 66 µM using Beer-lambert law:

$$A = \varepsilon cl$$

Where A is the absorbance at the excitation maximum (510 nm), e is the extinction coefficient in (M-1 cm-1 ), l is the pathlength in cm and C is the Molar concentration.
